# Gene expression profiling of lymph node sub-capsular sinus macrophages in cancer

**DOI:** 10.1101/2021.01.14.425337

**Authors:** Danilo Pellin, Natalie Claudio, Ferdinando Pucci

## Abstract

Lymph nodes are key lymphoid organs collecting lymph fluid and migratory cells from the tissue area they survey. When cancerous cells arise within a tissue, the sentinel lymph node is the first immunological organ to mount an immune response. Sub-capsular sinus macrophages (SSMs) are specialized macrophages residing in the lymph nodes that play important roles as gatekeepers against particulate antigenic material. In the context of cancer, SSMs capture tumor-derived extracellular vesicles (tEVs), a form of particulate antigen released in high amounts by tumor cells. We have recently demonstrated that SSMs possess anti-tumor activity because in their absence tumors grow faster. A comprehensive profiling of SSMs represents an important first step to identify the cellular and molecular mechanisms responsible for SSM anti-tumor activity. Unfortunately, the isolation of SSMs for molecular analyses is very challenging. Here, we combined an optimized dissociation protocol, careful marker selection and stringent gating strategies to highly purify SSMs. We provide evidence of decreased T and B cell contamination, which allowed us to reveal the gene expression profile of this elusive macrophage subset. Squamous cell carcinomas induced an increase in the expression of Fc receptors, lysosomal and proteasomal enzymes in SSMs. These results suggest that SSMs may be able to capture immune complexes for antigen processing and presentation to B and T cells on both MHC class I and II.

## Introduction

Lymph nodes are key lymphoid organs collecting lymph fluid and migratory cells from the tissue area they survey. When cancerous cells arise within a tissue, the sentinel lymph node is arguably the first immunological organ to mount an immune response. Within a lymph node, immune cells are highly organized into different anatomical compartments, and such architecture underlies its function (Qi et al. 2014). Afferent lymphatic vessels deliver lymph-bound antigens and migratory cells into the lymph node sub-capsular sinus. Here, a mesh of cells, mainly macrophages, filter lymph-bound antigens according to their size. Small, soluble antigens seep through lymphatic conduits and are channeled toward lymph node resident dendritic cells. Large and particulate antigens (>5nm in hydrodynamic radius) are retained by sub-capsular sinus macrophages (SSMs) (Gretz et al. 2000; Sixt et al. 2005). These specialized macrophages play important roles as gatekeepers against invading pathogens and in relaying immune complexed antigens to B cells for deposition on follicular dendritic cells (Phan et al. 2009; Gaya et al. 2015; Junt et al. 2007; Iannacone et al. 2010). In the context of cancer, SSMs capture tumor-derived extracellular vesicles (tEVs), a form of particulate antigen released in high amounts by tumor cells and overflowing into sentinel lymph nodes very early during disease progression (Pucci et al. 2016).

We have recently demonstrated that SSMs possess anti-tumor activity because in their absence tumors grow faster (Pucci et al. 2016). These observations have been confirmed in several tumor types and by other investigators (Louie 2019). However, the cellular and molecular mechanisms responsible for SSM anti-tumor activity are still unknown. In addition, we unveiled a link between tEVs, SSMs and immunoglobulins. Still, many details about SSMs’ contribution to the generation of humoral immune responses against tumor antigens is unclear. How do SSMs capture tEVs? Which signals SSMs release after capturing tEVs? How do they process tEV-bound antigens? A comprehensive profiling of SSMs represents an important first step to answer the above questions and to start elucidating their anti-tumor activity. Unfortunately, the isolation of SSMs for molecular analyses is very challenging. Likely due to their role as lymph node gatekeepers, SSMs are highly dendritic in shape and constantly extend pseudopods surveying the sub-capsular space (unpublished observations). These sub-cellular structures break apart during conventional enzymatic dissociation of lymph nodes and end up sticking to other cells, mainly lymphocytes (Gray et al. 2012). Indeed, initial attempts to profile SSMs showed a relatively high T and B cell contamination (Phan et al. 2009).

Here, we combined an optimized dissociation protocol, careful marker selection and stringent gating strategies to highly purify SSMs. We provide evidence of decreased T and B cell contamination, which allowed us to reveal the gene expression profile of lymph node macrophage subsets.

## Results

SSMs can be distinguished by other lymph node macrophage populations based on surface markers. Work from Dr. Cyster group showed that SSMs possess low phagocytic activity (Phan et al. 2009), which suggests that SSMs may display low side scatter during flow cytometry-based analyses. Our recent work confirmed this hypothesis (Pucci et al. 2016) and in this study we took advantage of this property to purify SSMs from tumor-draining lymph nodes (tdLNs) as 7AAD– CD45+ CD3– B220– Ly6G– CD11b+ CD169+ CD11c– SSC^LO^ (Figure 1A). As reference population, we isolated medullary sinus macrophages (MSMs) as 7AAD– CD45+ CD3– B220– Ly6G– CD11b+ CD169+ CD11c+ SSC^HI^. Non-draining lymph nodes (ndLNs) were also collected from the contralateral flank of mice carrying a squamous cell carcinoma model (MOC2), for a total of 4 macrophage subsets. SSMs and MSMs gated as above had physical scatter parameters consistent with a macrophagic origin (Figure 1B). We then performed deep sequencing on total RNA extracted from sorted samples (∼10 thousand cells each).

**Figure 1.**
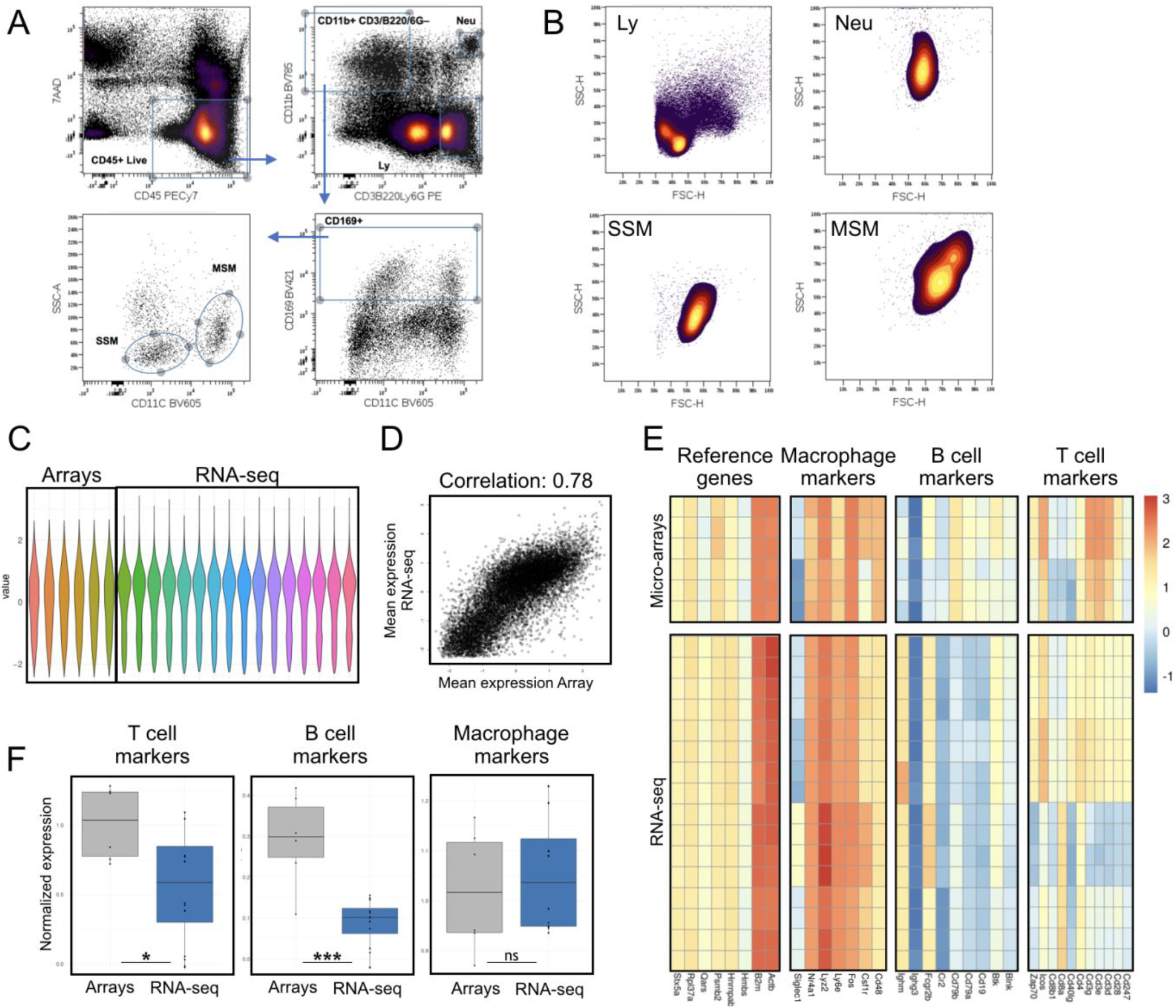
Comparison between previously published SSM gene expression arrays and our RNA sequencing data. A: Sorting strategy. B: Forward and side scatter parameters of SSM, MSM, lymphocytes (Ly, gated as in A) and neutrophils (Neu, gated as in A). C: Normalization of gene expression arrays and sequencing data for comparison. D: Normalization preserves high correlation between datasets (Pearson). E-F: Evaluation of T, B cell and macrophage markers, along with reference genes, shows significant reduction (F) of both T and B cell contamination in our sequencing data, as compared to previously published array data. Heatmaps show raw data normalized as in C.

In order to assess potential T and B cell contamination, we defined 4 gene signatures, one for macrophages, one for B cells, one for T cells and one consisting of reference genes. Before comparing these 4 gene signatures between our gene expression profiling data and previously published microarray data (GSE15767), we normalized the dataset to allow inter-platform comparison (Figure 1C-D). We observed that the reference genes and the macrophage signatures remained constant between the datasets, whereas the T and B cell signature was significantly lower (p=0.0266 and p=0.0008, respectively) in our dataset (Figure 1E-F). These results indicate that the isolation procedure adopted decreased T and B cell contamination in our lymph node macrophage preparations and that the resulting gene expression data is suitable for investigating SSM biology at the molecular level.

To start identifying SSM-specific markers, versus MSMs, we compared these two subsets in ndLNs. These analyses allowed us to highlight several gene families of interest, including general macrophage markers, immunoglobulins Fc receptors, integrins, C-type lectin receptors, cytokines and chemokines (Figure 2). As expected, SSMs expressed lower levels of the defining marker CD11c (Itgax).

**Figure 2.**
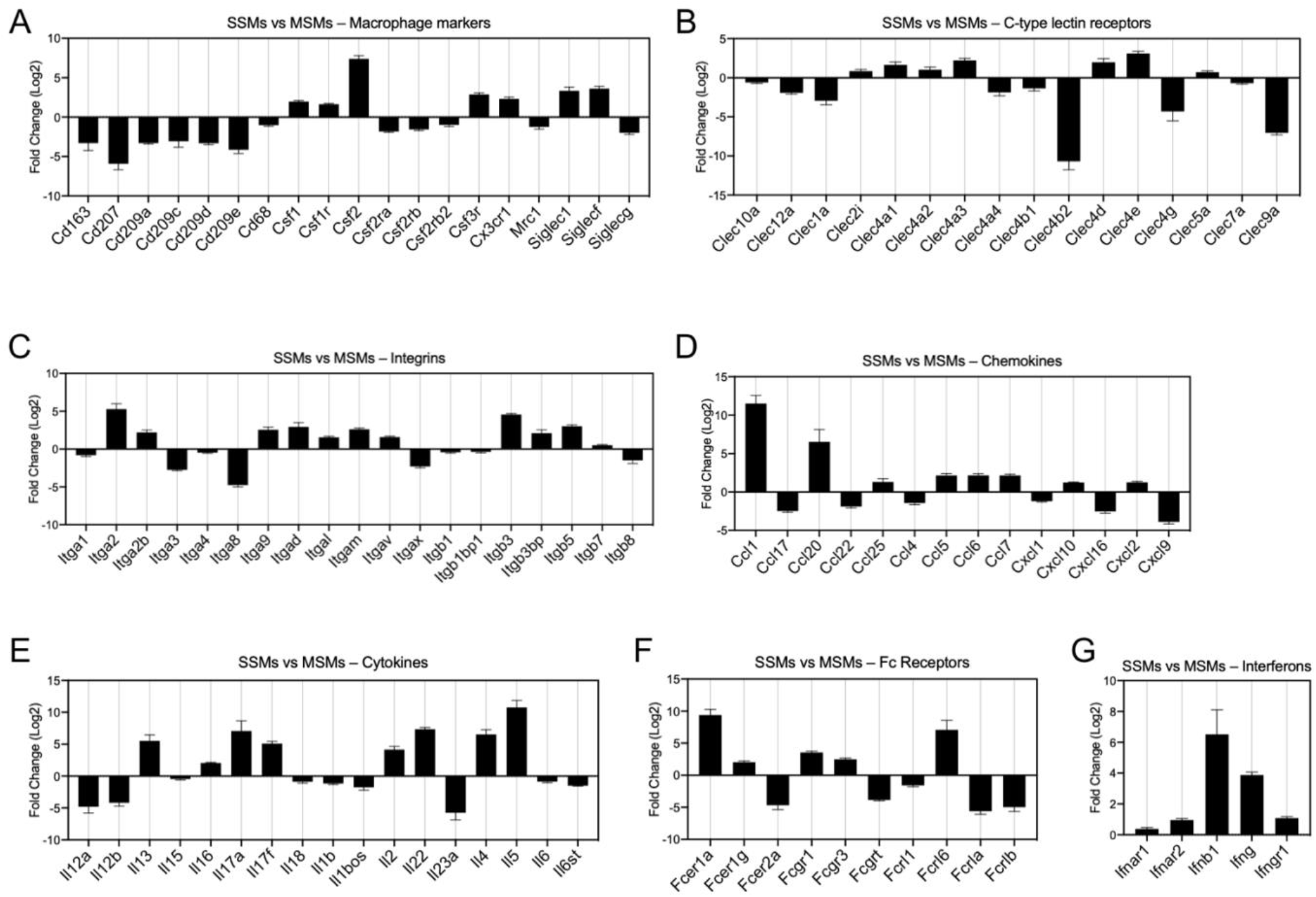
Comparison between SSMs and MSMs from non-draining lymph nodes. Genes shown have a p-adjusted value < 0.01. A: Macrophage markers. B: C-type lectin receptors. C: Integrins. D: Chemokines. E: Cytokines. F: Fc receptors. G: Interferons and receptors.

We next asked how the presence of a tumor influences SSMs gene expression. To this end, we compared SSMs profiles between tumor-draining and non-draining lymph nodes and performed gene ontology analysis on up-regulated and down-regulated genes. The results show which biological processes are affected by the presence of squamous cell carcinoma in the LN draining tissue. In particular, we observed an increase in expression of genes regulating immune responses and vesicular transport (Figure 3A-C). On the other hand, we observed a decrease in expression of genes involved in cell adhesion and T cell regulation (Figure 3D-E). SSMs displayed twice as many differentially expressed genes as compared to MSMs (Figure 3F), suggesting that tumors influence SSMs more than MSMs.

**Figure 3.**
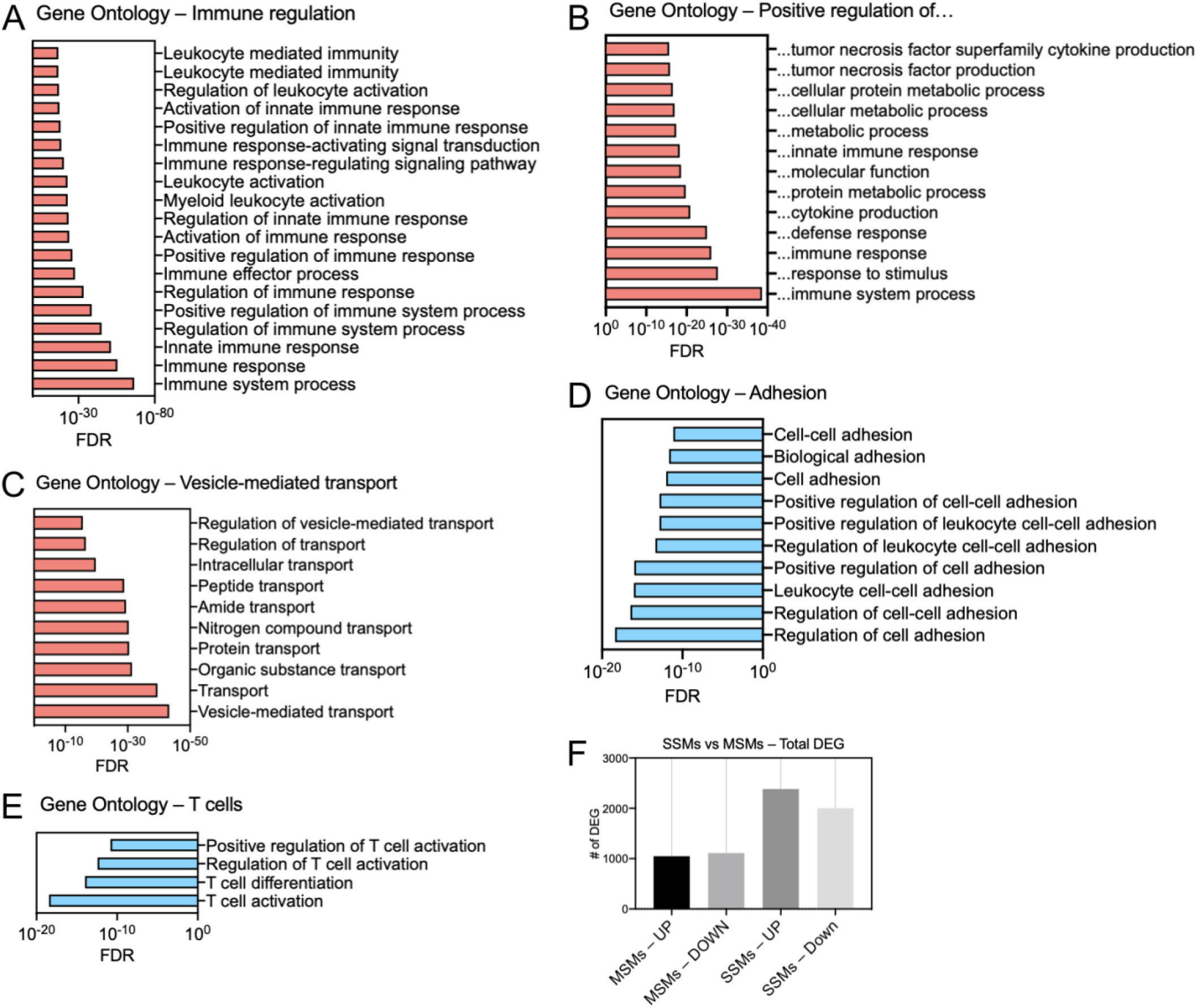
Gene ontology analysis of differentially expressed genes (DEG) in SSMs between tumor-draining and non-draining lymph nodes. A-E: Ontologies with upregulated genes are shown in red, ontologies with down-regulated genes in blue. Genes with a p-adjusted value < 0.01 and a mean expression > 100 were used. F: Total number of DEG in the two macrophage subsets between tumor-draining and non-draining lymph nodes.

To identify which genes in SSMs are influenced by a distant tumor, we compared SSMs profiles between tumor-draining and non-draining lymph nodes and focused on gene lists of interests (Figure 4). We started from gene lists from the ImmPort project (Bhattacharya et al. 2018), which included cytokines (chemokines, interferons and interleukins), TNF- and TGFb-family members, genes involved in antigen processing and presentation, and genes involved in innate immunity. Since it was reported that at steady state SSMs have low expression of lysosomal genes, we also analyzed Catepsins, vacuolar ATPases and other lysosome-associated genes (Figure 4C).

**Figure 4.**
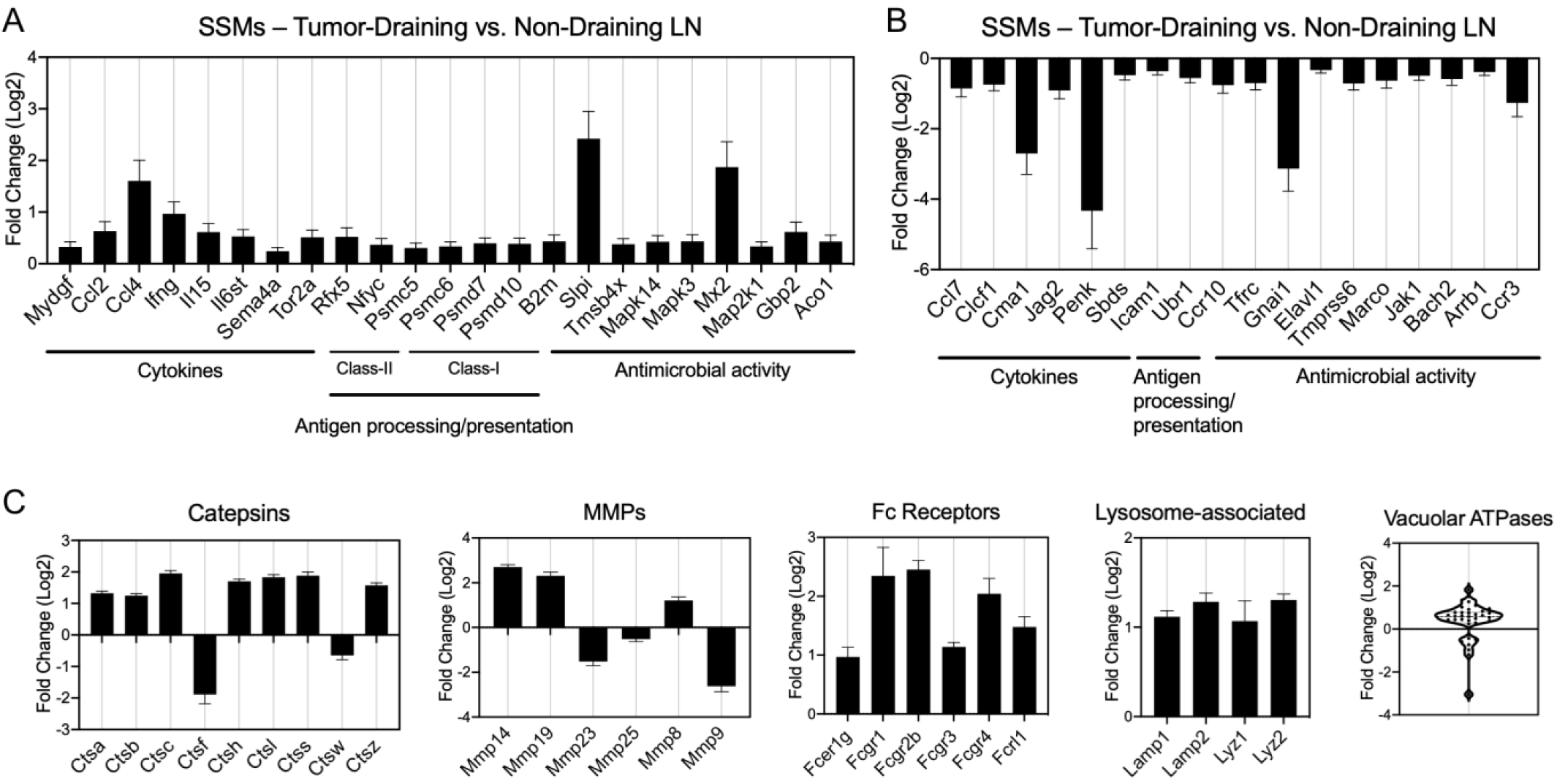
Gene families differentially regulated by tumors in SSMs. A-B: Immune gene lists, including cytokines, proteasomal factors and innate immune genes. C: Curated gene families, including lysosomal genes (Catepsins, vacuolar ATPases and other lysosome-associated genes), Fc receptors and matrix metalloproteases (MMPs). All comparisons are between SSMs from tumor-draining lymph nodes versus SSMs from non-draining lymph nodes. Genes with a p-adjusted value < 0.01 and a mean expression > 100 were used.

## Discussion

Sub-capsular sinus macrophages (SSMs) are known to be sensitive to enzymatic dissociation and tend to tear apart during isolation procedures (Gray et al. 2012). This issue makes molecular analyses challenging because SSM blebs stick to other lymphocytes generating a flow cytometric event with mixed macrophagic and lymphocytic markers. By coupling a short and gentle enzymatic dissociation protocol with a stringent gating strategy, we minimized T and B cell contamination in our SSM preparations. We employed a different combination of selection markers for flow sorting (i.e. gating both CD169^HI^ and CD169^LO^, and using CD11c/SSC as defining markers instead of F4/80 alone), which provided a better separation of the two lymph node macrophage subsets. These advancements allowed to get a first insight into the molecular biology of SSMs. Considering genes expressed at detectable levels (mean read count > 100), SSMs expressed higher levels of the following soluble factors, as compared to MSMs (Figure 2):

- CSF1 (M-CSF) and CSF2 (GM-CSF), two myelopoietic growth factors
- CCL1, involved in monocyte recruitment and endothelial cell migration
- IFNg, a key factor in anti-tumor immunity
- IL4, known to signal to B cells and recruited monocytes
- IL5, known to signal to eosinophils
- IL17a, IL17f and IL22, members of the T_H17_ cytokine family

These observations suggest a potential role for SSMs in the above processes. IL4 and IL5, although among the most highly DEG, are expressed at relatively low levels (∼200 raw counts). The T_H17_ cytokine family signature is of some concern since it was previously reported that SSM fragments are found on IL7Ra+ IL17-committed innate-like lymphoid cells (Gray et al. 2012). Of interest, SSMs expressed higher levels of Fcgr1 and Fcgr3, two Fc receptors known to bind activating immunoglobulins (Bournazos and Ravetch 2017). This last observation is consistent with the reported anti-tumor activity of these specialized macrophages (Pucci et al. 2016).

In the context of cancer, SSM gene profile was affected twice as much as compared to MSM gene profile (Figure 3F). Given that the sub-capsular sinus is the location where tEVs are first encountered, these observations further support the fact that the LN macrophage populations isolated represent bona-fide subsets residing in the sub-capsular and medullary sinuses, respectively. Squamous cell carcinomas induced an increase in immune regulatory gene networks and vesicular transport in SSMs. This observation, along with previously published functional data (Pucci et al. 2016), suggest that SSMs play key roles in the immune response to tumors. Interestingly, adhesion processes were down-regulated by tumors, potentially providing a mechanistic basis of previous observations indicating that SSM density is reduced in certain pathological conditions (Pucci et al. 2016; Gaya et al. 2015).

Squamous cell carcinomas induced an increase in the expression of several Fc receptors within SSMs, along with lysosomal and proteasomal enzymes (Figure 4). These results suggest that, in the context of cancer, SSMs may be able to capture immune complexes for antigen processing and presentation to B and T cells on both MHC class I and II.

Although these results represent a first step toward a better understanding of the biology of SSMs in cancer and beyond, more work is needed to validate the targets identified here, either as novel biomarkers of LN macrophage subsets or from a functional perspective. Moreover, a cross-species comparison with SSMs from patient sentinel nodes is still lacking.

## Methods

### Tumor models

The chemically-induced squamous cell carcinoma model MOC2 was purchased from Kerafast. For tumor formation, 5•10^5^ cells in 50ul of PBS were implanted in the flank of C57B/6 male mice (Charles River Labs) intradermally, near the inguinal lymph node, as previously described. After 2 weeks, we selected mice with similar tumor size for tdLN and ndLN collection. All procedures were in accordance with OHSU IACUC.

### Enzymatic dissociation of lymph nodes

LNs from 2 different mice were pooled: 2 tdLN and 6 ndLN (contralateral inguinal, axillary, brachial). Dissociation buffer was prepared by dissolving 3mg/ml of Collagenase IV (Worthington), 0.1 U/ml of DNase I (Roche), 2% FBS, pen/strep and L-glutamine in IMDM. Whole LNs were incubated in 5ml of dissociation buffer at 37°C and 225rpm agitation. After 15 minutes, LNs were mechanically dissociated by passing them through an 18G needle syringe at 2ml/second (LNs burst immediately) and put back at 37°C and 225rpm agitation. After 15 more minutes, the LN cell suspensions were mechanically dissociated by passing them through an 27G needle syringe at 0.5ml/second for 10 times. The LN cell suspension was then filtered on a 100um cell strainer and washed in 50ml of MACS buffer (2mM EDTA, 2.5% BSA in PBS). All centrifugation steps were performed at 500g for 10 minutes and supernatants were checked at the microscope for absence of cells. If significant cell numbers were observed (>0.1•10^6^/ml), supernatants were diluted 1:1 in MACS buffer and centrifuged again. LN cells were resuspended at ∼10^8^/ml for staining (∼1.5ml) in MACS buffer.

### Flow sorting and RNA extraction

All antibodies (and Fc Block) were from Biolegend, and were used at 2ul per 100ul of cell suspension, with the exception of CD45-PC7 and Ly6G-PE, which were used at 1ul per 100ul of cell suspension. Single stain controls were made by pooling a small amount of all samples. Flow sorting was done on a BD Aria and recovered ∼10,000 LN macrophages from each group. LN macrophages were then pelleted and lysed in RLT with B-mercapto-ethanol for RNA extraction (RNeasy micro kit), with on column DNAse treatment.

### RNA sequencing

RNA integrity was determined using the 2100 Bioanalyzer (Agilent). RNA was converted to cDNA with the SMART-Seq v4 Ultra Low Input RNA kit (Takara) and converted into sequencing libraries using the DNA Nano Kit (Illumina). Libraries were profiled on the 4200 TapeStation (Agilent) and quantified using the Kapa Library Quantification Kit (Roche) on a StepOnePlus Real Time PCR Workstation (Thermo). Sequencing was done on a HiSeq 2500 (Illumina). Fastq files were assembled using Bcl2Fastq2 (Illumina) and aligned using STAR aligner (Dobin et al. 2013).

### Gene expression analysis

Gene expression analysis was performed on 16 samples. The sample matrix is reported in Suppl. Table 1. We analyzed 4 pooled samples (biological replicates), each of them obtained from collecting cells from 2 mice. Every pool has generated 4 samples: CD11c+ (Phenotype=+) and CD11c-(Phenotype=-) cells sorted from Tumor Draining (TD=1) and Non-Draining (TD=0) lymph nodes. Raw sequencing files have been QC and aligned to the reference mouse genome mm10 using STAR workflow (Dobin et al. 2013), obtaining, as a result, the gene-based count matrices. We removed from the data set genes having a cumulative read count smaller than 10 and those that have been detected (reads count>0) in less than 2 samples, reducing the gene list from the initial 55423 to 20351. Statistical analysis has been performed in R (R Core Team 2013), using DEseq2 package (Love, Huber, and Anders 2014). Gene counts have been modeled using a negative binomial model, hypotheses tested by likelihood ratio test (LRT) and all p-value have been adjusted for multiplicity using Holm method (Holm 1979). A description of the model comparisons that have been performed and their motivation follows.

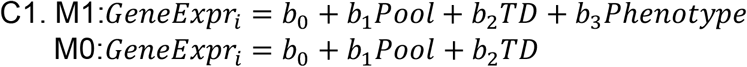

This comparison is focused on the identification of genes with a consistent expression modification between phenotypes, taking into account the possible confounding of TD and Pool factors. The complete list of genes is available in Suppl. Table 2.

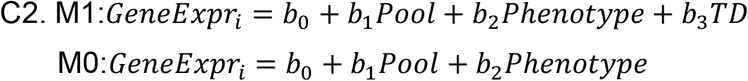

With this analysis, we investigated which are the genes with a significant association with TD variable, adjusting for Pool and Phenotype effects. The complete list of genes is available in Suppl. Table 3.

### Quantification of B- and T-cells contamination

We quantitatively assessed the B/T-cell fractions in our samples and compared them to previously published data (Phan et al. 2009) accessible through GEO code GSE15767. RNA-seq and microarray are platforms aimed at measuring gene activity level by mRNA quantification, but they rely on profoundly different techniques and protocols. In order to make the cross-platforms comparison as robust, meaningful, and accurate as possible, we implemented the following analysis workflow. Microarray technology uses several probes distributed along each gene’s mRNA sequence to have a reliable estimate of its expression; hence, we grouped and averaged the expression values based on gene identifier. We filtered out all genes detected only in one platform, reducing the gene list from 55423 to 14003. We normalized the log2 expression values in each sample in the 2 datasets by standardization (subtract mean and divide by standard deviation) and calculate platform-wise gene-specific expression levels by averaging all RNA-seq and microarrays data points. Based on the visual inspection of the scatter-plot in Fig. 1C and on Pearson correlation in Fig. 1D (r=0.78), we concluded that the two datasets are comparable. We defined shortlists of lineage-specific marker genes based on ImmPort gene lists (Bhattacharya et al. 2018):

- B-cell markers: Blnk, Btk, Cd19, Cd79a, Cd79b, Cr2, Fcgr2b, Ighg3, Ighm.
- T-cell markers: Cd247, Cd28, Cd3d, Cd3e, Cd3g, Cd4, Cd40lg, Cd8a, Cd8b1, Icos, Zap70.
- House Keeping: Rpl37a, B2m, Hmbs, Stx5a, Psmb2, Hnrnpab, Actb, Qars.
- Macrophages markers: Csf1r, Cd45, Siglec1, Cd48, Lyz2, Fos, Nr4a1, H2-k1, Ly6e.

For each sample, we calculated the lineage signature values by summing genes’ expression levels in the corresponding list (Fig 1E). Results are reported in Suppl Table 4. Housekeeping signature has low variability and is used as a reference to determine other lineages contribution. Boxplots of ratios are visible in Fig 1F for B/T-cell and Macrophages. Differences in ratio distributions tested using the non-parametric Mann-Whitney test.

